# The polyketide pathway in sporopollenin biosynthesis is specific to land plants (Embryophyta)

**DOI:** 10.1101/2024.10.10.617703

**Authors:** Damanpreet K. Sraan, Neil W. Ashton, Dae-Yeon Suh

## Abstract

**Background and Aims:** Sporopollenin (SP) is a complex biopolymer in the outer wall of spores and pollen and provides protection from environmental stresses. Its extraordinary chemical resistance, especially to acetolysis, was widely used to identify SP in biological specimens. This broad definition of SP led to claims for its widespread occurrence among diverse embryophyte and non-embryophyte taxa. We previously proposed a biochemical definition that can be used to distinguish genuine SP from other chemically resistant cell wall materials. The definition was centred on ASCL (Anther-Specific Chalcone synthase-Like), an embryophyte-specific enzyme of the polyketide pathway that provides precursors for SP biosynthesis. Herein, we examine the evolution and distribution of all five enzymes (CYP703A, CYP704B, ACOS, ASCL and TKPR) of the polyketide pathway and propose a new, more comprehensive definition of SP.

**Methods:** We performed BLASTp searches, phylogenetic tree construction, protein modeling and sequence analysis to determine the presence or absence of ACOS and TKPR in embryophytes and streptophytic algae.

**Key Results:** We found evidence that all five enzymes of the polyketide pathway evolved from ancestral enzymes of primary metabolism and ACOS, ASCL and TKPR were co-selected during evolution. The dosage of all five genes has been subjected to strict evolutionary control and, in some taxa, synteny has provided a selective advantage. All five enzymes are present in embryophytes but absent in green algae, indicating that the polyketide pathway and therefore SP is embryophyte-specific.

**Conclusions:** The addition of the polyketide pathway in the definition of genuine SP will allow separation of SP from algaenans and other chemically resistant ‘SP-like’ algal spore wall substances. This study further signifies SP as an evolutionary innovation unique to the embryophyte lineage and encourages research on possible evolutionary relationship between algal spore wall ‘SP-like materials’ and embryophyte SP.

## INTRODUCTION

### We previously proposed a new definition of sporopollenin (SP) as follows

“Sporopollenin is a chemically resistant complex heteropolymer present in the outer walls of spores and pollen grains and is composed partly of hydroxylated polyketides derived from the conserved polyketide pathway, which involves ASCL.” (Suh and Ashton, 2022).

As a protectant of progeny, SP is extraordinarily resistant to biological and chemical treatments, and it has been challenging to determine its chemical composition and structure. Hence, prior to our definition, resistance to acetolysis had been the sole criterion for the identification of SP (Rowley and Southworth, 1967; Heslop-Harrison, 1969). This imprecise definition resulted in claims for the existence of SP in algae and various microorganisms and also misguided research on the evolution of SP and its roles during terrestrialisation (Delwiche *et al*., 1989; Feofilova, 2010; de Vries and Archibald, 2018).

Recently, genetic and biochemical research has uncovered the biosynthesis pathways for various SP precursors (Grienenberger and Quilichini, 2021; Xue *et al*., 2023). The polyketide pathway comprised of five enzymes, CYP703A, CYP704B, ACOS (acyl-CoA synthetase), ASCL (anther-specific chalcone synthase-like) and TKPR (tetraketide α-pyrone reductase), provides polyketide SP precursors and is conserved in embryophytes. ASCL belongs to the type III polyketide synthase family, and previous phylogenetic studies of the family indicated that ASCL is specific to embryophytes (Jiang *et al*., 2008; Shimizu *et al*., 2017). Indeed, we discovered at least one *ASCL* in representative embryophytes from all lineages that produce spores or pollen with sporopolleninous exine. In sharp contrast, we failed to find any *ASCL* in the genomes of all algal species for which data are available, including Charophytes that were considered to produce SP-like substances. This finding led us to propose the above-mentioned definition for SP to distinguish genuine SP from other similarly resistant substances and to use *ASCL* as an identifying marker for the presence of SP in spore and pollen walls. We recognised that there are other enzymes than ASCL in the polyketide pathway and exploring algal genomes for orthologues encoding these enzymes had the potential for augmenting our SP definition (Suh and Ashton, 2022).

In this study, we investigated the evolution of the genes in the polyketide pathway and examined whether CYP703A, CYP704B, ACOS and TKPR are conserved in embryophytes and whether they are absent in green algae.

## METHODS AND MATERIALS

### Database searches and phylogenetic analysis

BLASTp searches were performed against 35 embryophytes selected from all major lineages from hornworts to eudicots in Phytozome 13 (https://phytozome-next.jgi.doe.gov/), the National Center for Biotechnology Information (https://www.ncbi.nlm.nih.gov/), the *Takakia* project (https://www.takakia.com/blast/blast_cs.html), the *Anthoceros* genomes (https://www.hornworts.uzh.ch/en/hornwort-genomes.html) and the FernBase (https://fernbase.org) using PpACOS6, AtACOS5, PpTKPR1 and AtTKPR1 as queries. Similarly, BLASTp searches were performed against each algal genome in PhycoCosm (https://phycocosm.jgi.doe.gov/phycocosm/home) (accessed on 12 March 2024) using the same query sequences. The cutoff E value was 1.0 × E^−20^ and truncated and redundant sequences were removed. Representative sequences of the 4CL (4-coumarate:CoA ligase) and CCR (cinnamoyl-CoA reductase) families were retrieved from Phytozome 13. The sequences used for phylogenetic tree construction (Supplementary Data Table S1, S2) were aligned by Clustal Omega (https://www.ebi.ac.uk/jdispatcher/msa/clustalo), and Maximum Likelihood (ML) trees were built in MEGA 11 under the WAG+G+I substitution model. Support for the tree was measured using 1,000 bootstrap replicates.

### Protein modeling and sequence analysis of ACOS and TKPR

Structures of ACOS from *Physcomitrium patens*, arabidopsis, *Oryza sativa* (PpACOS6, AtACOS5, OsACOS12, respectively) and that of *P. patens* TKPR (PpTKPR1) were modeled with SWISS-MODEL (https://swissmodel.expasy.org/) and AlphaFold (https://alphafold.ebi.ac.uk/). The modeled structures of ACOSs were compared to the X-ray crystal structure of tobacco 4CL2 (Nt4CL2; PDB id, 5BST) and their active site sequences were aligned using Clustal Omega to reveal enzyme-specific residues. Likewise, the modeled structure of PpTKPR1 was compared to crystal structures of *Medicago truncatula* CCR (MrCCR; PDB id, 4R1U), *Panicum virgatum* dihydroflavonol 4-reductase (PvDFR; PDB id, 8FEM) and *Sorghum bicolor* anthocyanidin reductase (SbANR; PDB id, 8FIP) and their active site sequences were aligned using Clustal Omega.

### Synteny analysis

Coordinates for genes of the polyketide pathway in the taxa discussed and total scaffold lengths of their genomes were obtained from Phytozome 13. The coordinates were used to calculate the lengths of DNA spanning chosen pairs of syntenic genes (from the beginning of one gene to the end of the other). Total genomic scaffold lengths were assumed to represent total genomic euchromatin. The ratio of these values was viewed as the probability by chance alone of one gene being present within the spanned DNA on a particular chromosome. The probability of n genes being located in this span of DNA was calculated as the ratio to the power, n. The uncorrected probabilities for *Arabidopsis thaliana* were calculated in this way using data from the arabidopsis TAIR 10 genome sequence in Phytozome 13. The values corrected for heterochromatin utilised data from the recently sequenced arabidopsis Col-XJTU assembly (Wang *et al*., 2022): the length of CEN1 (3.8 Mb) was subtracted from the length of DNA spanned by the five syntenic genes of the polyketide pathway (26 Mb) located on Chr 1 and the combined length of the five CENs (21.7 Mb) and two NORs (0.64 Mb) was subtracted from the genome size (134 Mb). This gave a length for euchromatin spanned by five syntenic genes of the pathway on Chr 1 of 22.2 Mb and for total genomic euchromatin of 112 Mb. Corrected probabilities of the observed synteny on Chr 1 and Chr 4 occurring by chance alone were then obtained by entering these calculated values in the equation given in the main text.

## RESULTS AND DISCUSSION

### The embryophyte-specific polyketide pathway and an expanded definition of sporopollenin

It had been suggested previously that ACOS and TKPR are embryophyte-specific (de Azevedo Souza *et al*., 2008; Grienenberger *et al*., 2010). However, information about algal homologues of these enzymes was lacking. Taking advantage of newly determined genome sequences, we expanded earlier studies and searched for ACOS and TKPR in the genomes of thirty-five embryophytes representing all major lineages (Supplementary Data Tables S1, S2). Putative ACOS sequences collated from BLASTp searches were identified based on phylogenetic clustering with known enzymes (Supplementary Data Figs S1, S2). In parallel, for additional support for enzyme identification, we searched for amino acid residues that are conserved in ACOS but not in its paralogues. ACOS and 4CL, both of which belong to the 4CL family, catalyze formation of CoA esters. Both enzymes use ATP and CoA as co-substrates, but they differ in their acid substrates. While ACOS accepts hydroxyfatty acids (provided by CYP703A and CYP704B in the polyketide pathway, Fig 1) as substrate, 4CL accepts 4-coumaric acid derivatives. Crystal structures of several 4CLs pinpointed active site residues responsible for substrate specificity of the enzyme (Hu *et al*., 2010). Sequence alignment of representative enzymes of the 4CL family (Supplementary Data Fig. S3) and comparison of 4CL crystal structures (e.g., tobacco 4CL2) with ACOS structure models (Fig. 2A, B) indicated that Gly^241^ and S^339^C(V/I)Tx(S/T) (numbering of AtACOS5) are uniquely conserved in ACOS and could serve as a determinant of ACOS. Reflecting the different substrate specificity of ASCL and 4CL, corresponding residues in Nt4CL2 are Ser^243^ and G^339^PVLxM (Fig. 2B). We then performed a BLASTp search of all algal genomes in the PhycoCosm database for algal homologues of ACOS (Supplementary Data Table S1) and constructed a phylogenetic tree with representative embryophyte enzymes of the 4CL family. As shown in Fig. S1, none of the algal homologues, including 13 from charophytes, is found in the ACOS clade and none of them possesses the above-mentioned ACOS signature residues (Supplementary Data Fig. S3). We concluded that none of the algae examined has an ACOS.

**Fig 1.**
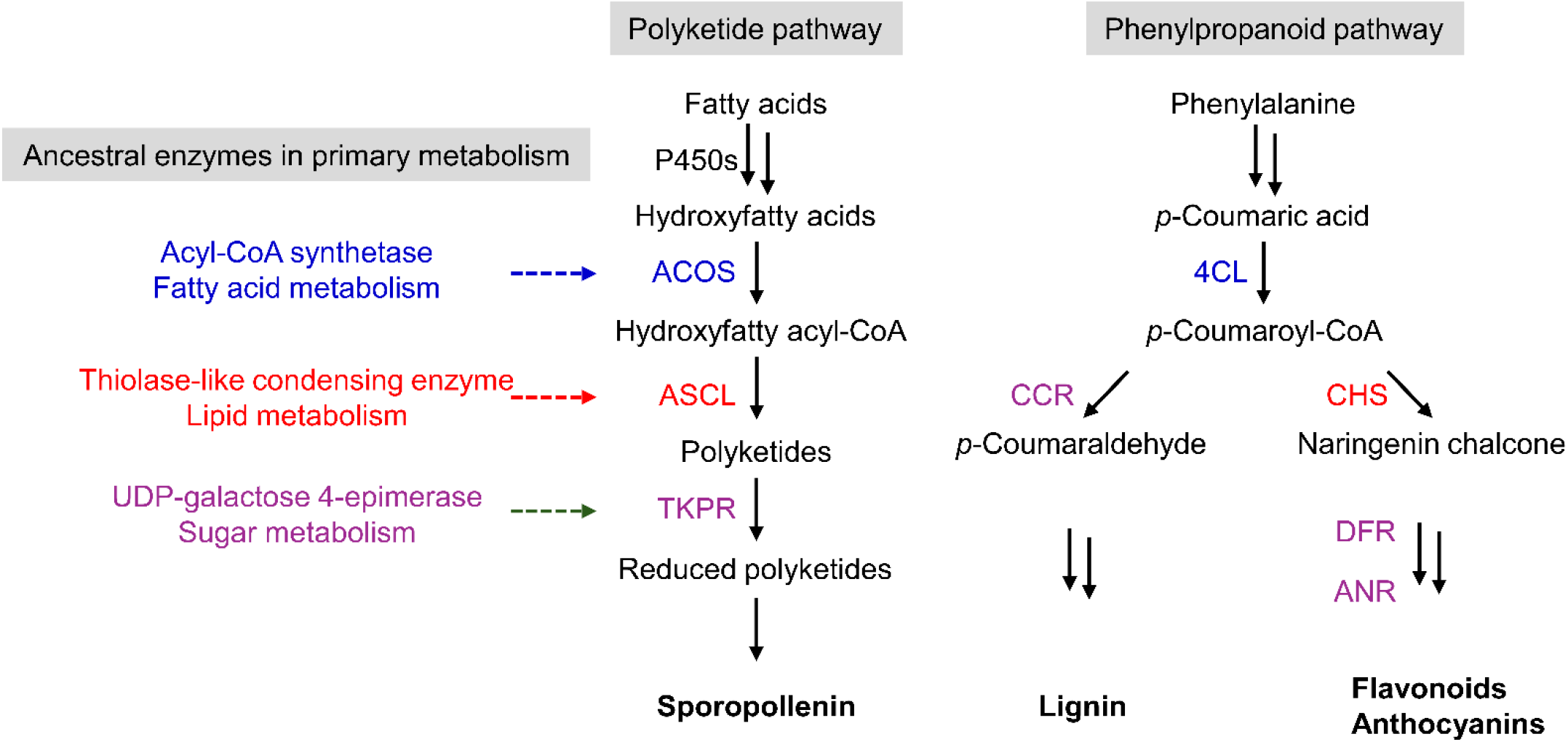
The origin and evolution of the polyketide pathway in sporopollenin biosynthesis. ACOS, ASCL and TKPR in the polyketide pathway are shown with their evolutionarily related enzymes in the phenylpropanoid pathway. Also shown are the enzymes of primary metabolism proposed to share the same ancestors with the related enzymes in the polyketide and phenylpropanoid pathways. Related enzymes are indicated by text in the same colour. For instance, ACOS and 4CL are thought to have evolved from the ancestral form of acyl-CoA synthetase in fatty acid metabolism and are presented using blue text. 4CL, 4-coumarate:CoA ligase; ACOS, acyl-CoA synthetase; ANR, anthocyanidin reductase; ASCL, anther-specific chalcone synthase-like; CCR, cinnamoyl-CoA reductase; CHS, chalcone synthase; DFR, dihydroflavonol 4-reductase. For further information about the enzymes in the pathways, readers are referred to Grienenberger and Quilichini (2021) and Ferrer et al. (2008).

**Fig 2.**
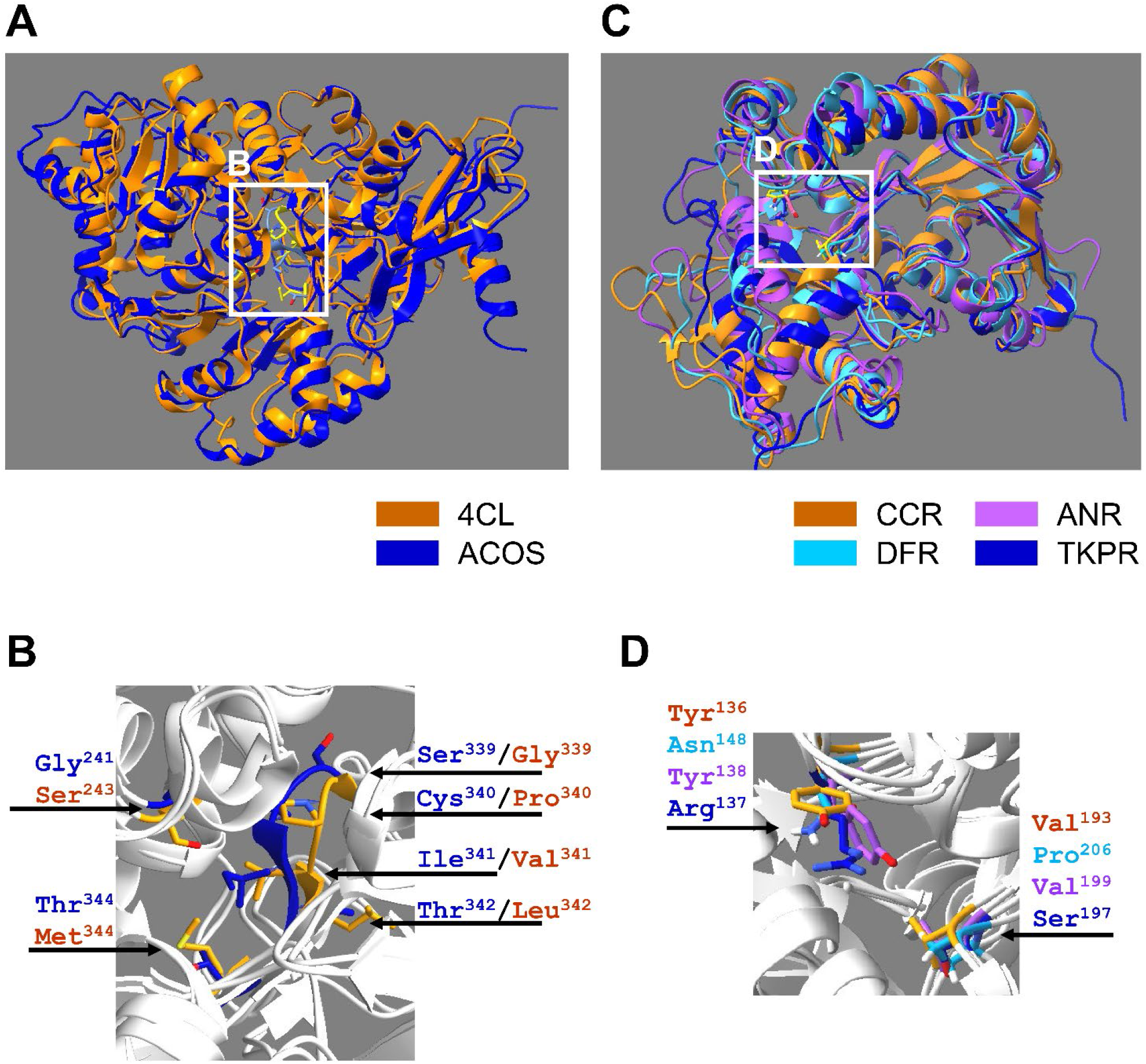
ACOS- and TKPR-specific residues at their substrate binding sites. (**A, B**) The modeled structure of arabidopsis ACOS (AtACOS5, blue) is superimposed with the crystal structure of tobacco 4CL (Nt4CL2, PDB id, 5BST, orange). Their acid substrate binding pockets (white-boxed) are magnified in **B** to show the uniquely conserved residues in ACOS (Gly^241^, Ser^339^, Cys^340^, Ile^341^, Thr^342^ and Thr^344^) and 4CL (Ser^243^, Gly^339^, Pro^340^, Val^341^, Leu^342^ and Met^344^). These substrate-determining residues, also shown in Supplementary Data Fig. S3, allow facile identification of ACOS from its paralogues, including 4CL. (**C, D**) The modeled structure of *P. patens TKPR* (PpTKPR1, blue) is superimposed with the crystal structures of *Medicago truncatula* CCR (MtCCR; PDB id, 4R1U, orange), *Panicum virgatum* DFR (PvDFR; PDB id, 8FEM, cyan) and *Sorghum bicolor* ANR (SbANR; PDB id, 8FIP, purple). Their substrate binding pockets (white-boxed) are magnified in **D** and the TKPR-specific Arg^137^ and Ser^197^ in PpTKPR1 are substituted with Tyr^123^ and Val^180^ in MtCCR, Asn^148^ and Val^199^ in PvDFR, and Tyr^138^ and Pro^206^ in SbANR (Supplementary Data Fig. S4), conferring their different substrate specificity.

Similarly, sequences and modeled structures of TKPRs were compared with those of paralogous DFR, ANR and CCR to identify two substrate binding site residues Arg^137^ and (Ser/Thr)^197^ (numbering of PpTKPR1) as determinants of TKPR (Supplementary Data Fig. S4; Fig. 2C, D). BLASTp searches yielded a total of 29 algal homologues, including four from charophytes (Supplementary Data Table S2). None of them resides in the TKPR clade (Supplementary Data Fig. S2) and, furthermore, no charophyte homologue has the TKPR signature residues, Arg and (Ser/Thr). The cytochrome P450 family had already been extensively studied and meticulously curated in the P450 database (https://drnelson.uthsc.edu/ , Nelson, 2009).

CYP703A is present only in embryophytes, while CYP704B is found in embryophytes, protists and red algae, but, importantly, not in green algae (Morant *et al*., 2007; Li *et al*., 2010; Hansen *et al*., 2021). These findings led us to conclude that all of the five enzymes in the polyketide pathway are present in embryophytes but absent in green algae, and produce polyketide precursors of genuine SP. Therefore, we have expanded our definition of SP as follows:

“Sporopollenin is a chemically resistant complex heteropolymer present in the outer walls of spores and pollen grains and is composed partly of hydroxylated polyketides derived from the conserved polyketide pathway, which typically involves CYP703A, CYP704B, ACOS, ASCL and TKPR.”

### Evolutionary origin of the polyketide pathway

Next we searched the literature for studies on the evolutionary origins of the enzymes in the polyketide pathway. It is generally accepted that enzymes of secondary metabolism have been derived from pre-existing enzymes in primary metabolism via gene duplication and neofunctionalisation (Firn and Jones, 2000).

#### ACOS

The 4CL family belongs to the adenylate-forming enzyme superfamily present in all organisms. The superfamily shares a common reaction of the formation of an adenylated intermediate as in fatty acid activation by fatty acyl-CoA synthetase. 4CL produces CoA esters of hydroxycinnamic acids which are intermediates in the biosynthesis of a vast array of phenolic secondary metabolites, including monolignols and flavonoids (Hahlbrock and Scheel, 1989). Based on kinetic and phylogenetic analysis, it has been suggested that the members of the superfamily share a common ancestor similar to extant acyl-CoA synthetase in primary fatty acid metabolism (Fig. 1) (Linne *et al*., 2007; de Azevedo Souza *et al*., 2008).

#### ASCL

ASCL belongs to the family of type III polyketide synthases (PKS). Type III PKSs condense a starter CoA substrate with a number of (mostly three) malonyl-CoA molecules and produce a variety of polyketide secondary metabolites with different biological functions (Abe and Morita, 2010). There are more than twenty type III PKSs in plants and the most well-known is chalcone synthase (CHS) that catalyzes the first committed step in flavonoid biosynthesis. Type III PKSs, along with several condensing enzymes in fatty acid metabolism, belong to the thiolase superfamily. An earlier phylogenetic study showed that type III PKSs have evolved, along with β-ketoacyl synthases in fatty acid synthesis, from an ancestral thiolase-like condensing enzyme in lipid metabolism (Fig. 1) (Jiang et al. 2008).

#### TKPR

TKPR and its paralogues DFR and CCR all belong to the short-chain dehydrogenase/reductase (SDR) superfamily present in all living organisms. DFR plays a role in the biosynthesis of anthocyanins while CCR catalyses the first committed step of lignin biosynthesis (Lacombe *et al*., 1997; Petit *et al*., 2007). Enzymes in the SDR family display similarities to sugar-metabolizing enzymes in bacteria and it has been suggested that an ancestor of the superfamily could have been UDP-galactose-4-epimerase in primary sugar metabolism (Fig. 1) (Baker and Blasco, 1992; Kallberg *et al*., 2002).

#### CYP703A and CYP704B

The CYP family is one of the largest enzyme families and present in all kingdoms of life. The family is made of many clans and each clan contains individual CYP enzymes. The CYP 71 and 86 clans, to which CYP 703A and 704B belong respectively, are mainly involved in fatty acid metabolism. All extant CYPs are monophyletic, and CYP51 (sterol 14α-demethylase) in sterol biosynthesis is widely accepted as the ancestral progenitor (Yoshida et al. 2000; Lamb et al., 2021). To summarize, it appears that ACOS, ASCL and TKPR all evolved from ancestral enzymes in primary metabolism in the common ancestor of embryophytes. It could be more than a coincidence that their paralogues play important roles in phenylpropanoid pathways (Fig. 1) and it may reflect critical contributions that SP and phenylpropanoids made during early evolution of land plants.

### ACOS, ASCL and TKPR were co-selected during evolution

Earlier, we found that the genome of *Zostera marina* (a seagrass) that produces exine-less pollen does not contain *ASCL*, which we attributed to secondary gene loss (Suh and Ashton, 2022). The *Z. marina* genome also lacks an *ACOS*, and it contains one copy of a *TKPR*-like sequence that would produce a truncated TKPR-like protein because of nonsense mutation and pseudogenisation. Also, *Zingiber officinale* produces pollen with a reduced exine with regular discontinuities (Subbarayudu *et al*., 2014). A tBLASTn search against the genome of *Z. officinale* Roscoe (taxid:94328) (https://blast.ncbi.nlm.nih.gov/, accessed in February 2024) failed to find *ACOS, ASCL* or *TKPR*. Therefore, in line with our new and expanded definition, neither *Z. marina* nor *Z. officinale* makes genuine SP. Based on these data, we also concluded that the three genes of the polyketide pathway were co-selected during embryophyte evolution; loss of one of these genes would have alleviated the selective pressure for retention of the remaining genes in the pathway. *CYP703A* and *CYP704B* genes were found in the *Z. marina* and *Z. officinale* genomes unlike the other three genes in the polyketide pathway. That they did not suffer gene loss is probably because the CYP enzymes they encode are required for the biosynthesis of cutin monomers in addition to SP biosynthesis (Li *et al*., 2010).

### Gene dosage: Copy number of the genes in the polyketide pathway

We examined whether the copy number of the five genes in the pathway is controlled. The *CYP* family underwent diversification bursts (Nelson, 2019) and clans 71 and 86 are the two largest clans in embryophytes. For instance, the genome of *P. patens* has 71 CYPs and 41 of them are in clan 71. Nonetheless, *CYP703A* is often the only single gene family in the *CYP71* clan (Nelson, 2019). Also, *CYP704* is often maintained as a single copy or at low copy numbers (Hansen *et al*., 2021). In the 4CL family, while bona fide *4CL* genes have undergone gene family expansion, *ACOS* genes are encoded by single-copy genes in most plants (de Azevedo Souza *et al*., 2009). Out of 35 plants we examined, only the genomes of *P. patens* and *Glycine max* have two copies of *ACOS* and the rest have a single copy (Supplementary Data Table S1). Likewise, whereas multiple copies of *CHS* genes (encoding chalcone synthases) are found in most plants, only one (bryophytes, lycophytes, gymnosperms) or two copies of *ASCL* (angiosperms) were found in plant species that we examined (Suh and Ashton, 2022). As for *TKPR*, we found two copies of *TKPR* in most plant species and a single copy in a few species including *Ginkgo biloba* and *Spirodela polyrhiza* (Supplementary Data Table S2). These data indicate that gene dosage of all five genes in the pathway has been subjected to strict control during evolution. Indeed, the genomes of arabidopsis and rice contain one copy each of *CYP703A, CYP704B* and *ACOS* and two copies each of *ASCL* and *TKPR*. The genome of *P. patens* has one copy each of *ACOS* and *ASCL* and two copies of *TKPR* despite undergoing two whole genome duplications (Rensing *et al*., 2008; Barker and Ashton, 2016; Lang *et al*., 2018). We argue that the evolutionary benefits of gene dosage control were maintenance of steady metabolic flux and avoidance of the accumulation of intermediate(s) that could be prematurely incorporated into SP, harming its structural integrity.

### Synteny of genes of the polyketide pathway in model plants

Synteny of genes of the polyketide pathway can be observed in some model plants for which sequenced whole genome assemblies are available. The most obvious example is arabidopsis in which five genes, *CYP703A2, PKSA, ACOS, TKPR2* and *CYP704B*, are located on Chromosome 1 (Chr 1) and two genes, *PKSB* and *TKPR1*, are present on Chr 4. (*PKSA* and *PKSB* are *ASCL* orthologues in arabidopsis (Kim *et al*., 2010)). It is unlikely that the synteny of Chr 4 resulted from a recent segmental duplication since *PKSB* and *TKPR1* on Chr 4 are much closer together (0.23 Mb ) than *PKSA* and *TKPR2* on Chr 1 (25.4 Mb) and are not separated by a recognisable *ACOS* gene or CEN4 (Fig. 3).

**Fig 3.**
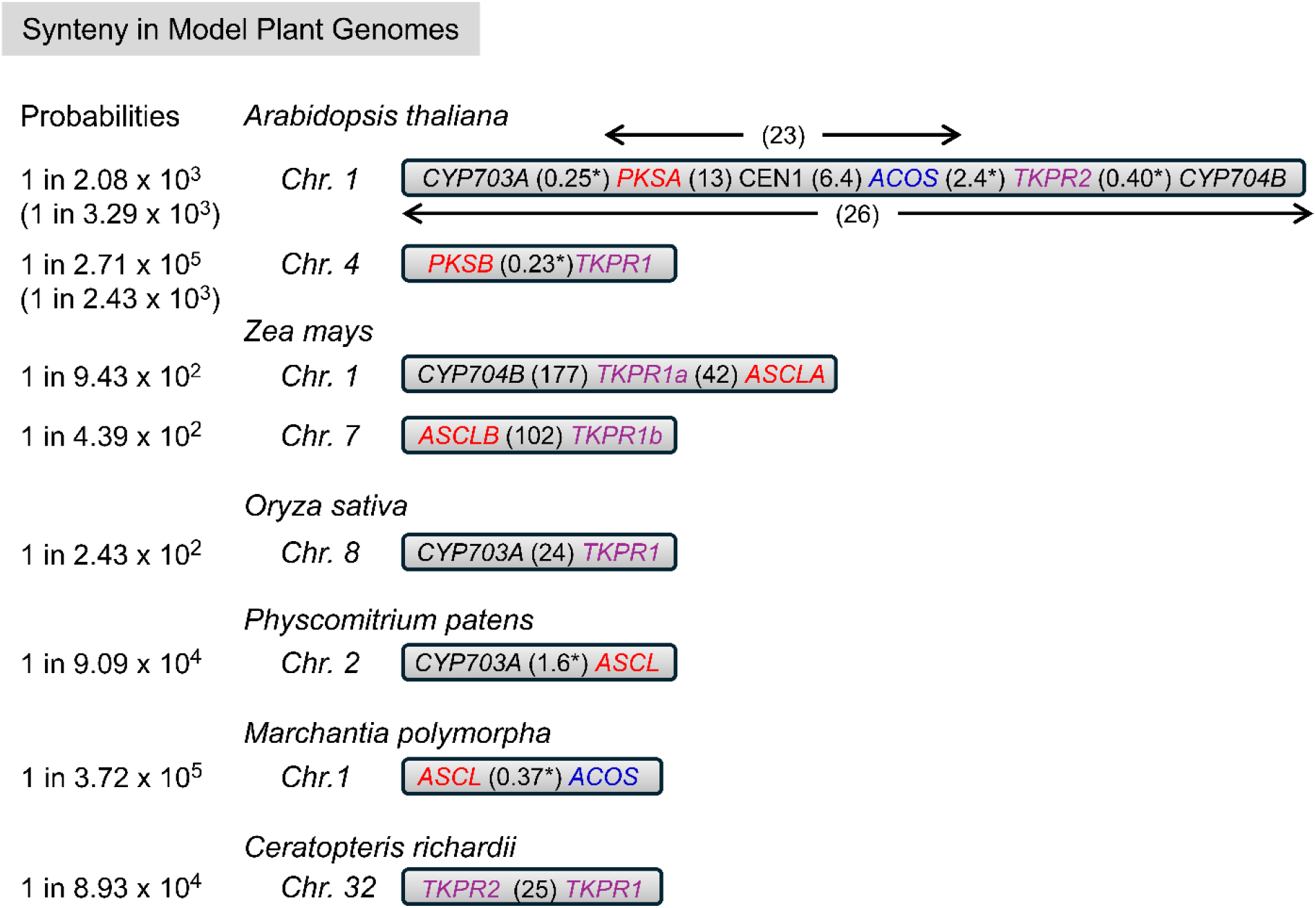
Synteny of genes of the polyketide pathway in model plant genomes The ideograms are not drawn to scale. Genes are colour-coded as follows: *CYP703A* and *CYP704B* (black), *ASCL* (red), *ACOS* (blue), *TKPR* (purple). (*PKSA* and *PKSB* are *ASCL* orthologues in arabidopsis). Numbers in brackets refer to DNA in Mb spanning pairs of genes. CEN1 (3.8 Mb) is approximately 15 Mb from the left end of Chr 1 of arabidopsis and separates *PKSA* and *ACOS*. Distances within or close to the range (0.2−2.0 Mb) for a euchromatin loop in the arabidopsis chromocenter-loop model of interphase chromosomes are indicated by an asterisk. Probabilities by chance alone of the observed degrees of synteny were calculated (*Prob* = *(length of DNA spanned by syntenic genes of the pathway in a single chromosome/total genomic scaffold length)*^*n*^. *Where n = number of syntenic genes in the chromosome*) and are shown as fractions given to three significant figures. The values shown in brackets for arabidopsis have been corrected for heterochromatin. Data used for this figure were obtained from Phytozome 13 and also, in the case of arabidopsis, from Wang *et al*. (2022).

We have calculated the probabilities by chance alone of the detected levels of synteny in arabidopsis, correcting for confounding factors by subtracting the amounts of heterochromatin (centromeres (CENs) and nucleolar organising regions (NORs)) in each chromosome and in the whole genome and calculating the actual span of syntenic genes in the euchromatin, as follows:

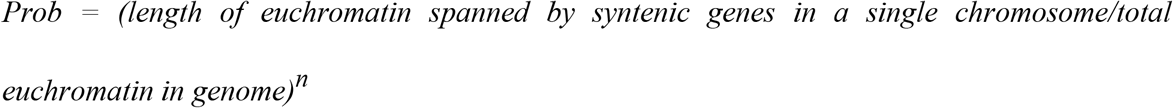

Where n = number of syntenic genes in the chromosome

Based on our calculations, the probabilities of the observed synteny arising by chance alone are low, e.g. 1 in 3,290 for the five genes on Chr 1 (This probability estimate may be higher than the actual probability since the CEN1 sequence contains a gap and, based on the physical map, CEN1 is approx. 9 Mb (Hosouchi *et al*., 2002)) and 1 in 243,000 for *PKSB* and *TKPR1* on Chr 4. We conclude that synteny must have provided a selective advantage during the evolution of SP. The most obvious possibility is that it facilitated epigenetic coordinate gene regulation thereby ensuring appropriate spatio-temporal expression. Interestingly, some syntenic genes, e.g. *PKSB* and *TKPR1* that span approx. 0.2 Mb of euchromatin on Chr 4, are close enough together to occupy a single euchromatin loop (0.2–2 Mb) in the chromocenter-loop model of arabidopsis interphase chromosomes (Fransz *et al*., 2002). That said, it is also clear that the number of syntenic genes of the pathway and their degree (closeness) of synteny varies widely between taxa (Fig. 3). Three genes of the polyketide pathway are located on Chr 1 and two more on Chr 7 of *Zea mays*, two are on Chr 8 of rice, two are on Chr 2 of *Physcomitrium patens*, two are on Chr 1 of *Marchantia polymorpha* and two are on Chr 32 of *Ceratopteris richardii*. Interestingly, a chromosome in each of four taxa, Chr 1 of arabidopsis, Chr 1 of *Z. mays*, Chr 8 of rice and Chr 2 of *P. patens* possesses a *CYP703A* or *CYP704B* gene plus either a *TKPR* gene or an *ASCL* orthologue or both. The distances between six gene pairs, four in arabidopsis, one in *P. patens* and one in *M. polymorpha* are such that each gene pair could be accommodated within a single euchromatin loop. However, we conclude other means of achieving coordinate gene expression must be operating as well as epigenetic mechanisms. Also we cannot rule out synteny conferring a selective advantage by an entirely different mechanism. As more sequenced genome level assemblies become available, it will be interesting to discover how widespread this phenomenon is.

## Conclusions

All evidence indicates that the polyketide pathway and therefore SP is embryophyte-specific. The addition of the polyketide pathway in the definition of genuine SP will allow SP to be distinguished unequivocally from algaenans and other chemically resistant ‘SP-like’ algal spore wall substances.

This new, expanded definition of SP implies that SP was a major evolutionary innovation unique to the embryophyte lineage that may have contributed to plant terrestrialisation. This study should also spur research on possible evolutionary relationships between genuine embryophyte SP and other ‘SP-like materials’ in reproductive bodies including algal zygospores and dinoflagellate cysts.

## SUPPLEMENTARY DATA

Supplementary data are available online at https://academic.oup.com/aob and consist of the following.

Table S1: Sequences used for 4CL family tree construction (Fig S1).

Table S2: Sequences used for CCR family tree construction (Fig S2).

Fig. S1: Maximum Likelihood tree of embryophyte ACOS and 4CL sequences and algal homologues.

Fig. S2: Maximum Likelihood tree of TKPR and other enzymes of the CCR family and algal homologues.

Fig. S3: Multiple sequence alignment of ACOSs, 4CLs and algal homologues.

Fig. S4: Multiple sequence alignment of representative enzymes of the embryophyte CCR family and algal homologues.

## FUNDING

This work was supported by the Natural Sciences and Engineering Research Council of Canada Discovery grant [RGPIN-2018-04286 to D.-Y.S.). D.K.S. was supported in part by University of Regina Graduate Scholarships.

## ACKNOWLDEGEMENTS

We thank William B. Browing (Southern Illinois Univ., USA) for informing us of the absence of *ASCL* in the *Zingiber officinale* genome. D-YS conceived the concept and D-YS, DKS and NWA collected and analysed data. D-YS and NWA wrote the manuscript.

## Notes

### Competing Interest Statement

The authors have declared no competing interest.

### Summary of Updates

A section on protein modeling for the identification of enzyme-specific residues is added. A figure (Fig 2) is added.

