## Supplementary Data for "The polyketide pathway in sporopollenin biosynthesis is specific to land plants (Embryophyta)"

The following Supplementary Data are available for this article:

Table S1: Sequences used for 4CL family tree construction (Fig S1).

Table S2: Sequences used for CCR family tree construction (Fig S2).

Fig. S1: Maximum Likelihood tree of embryophyte ACOS and 4CL sequences and algal homologues.

Fig. S2: Maximum Likelihood tree of TKPR and other enzymes of the CCR family and algal homologues.

Fig. S3: Multiple sequence alignment of ACOSs, 4CLs and algal homologues.

Fig. S4: Multiple sequence alignment of representative enzymes of the embryophyte CCR family and algal homologues.

**Table S1 Sequences used for 4CL family tree construction (Fig S1)**

| Name | Gene ID | Species | Classification |
|---|---|---|---|
| **Algal homologues** | | | |
| Gs233 | Galsul1\|233 | *Galdieria sulphuraria* | Rhodophyta, Galdieriales |
| Gs5225 | Galsul1\|5225 | *Galdieria sulphuraria* | Rhodophyta, Galdieriales |
| Gd7747 | Gradom1\|7747 | *Gracilaria domingensis* | Rhodophyta, Gracilariales |
| Pp5353 | Porpu1328_1\|5353 | *Porphyridium purpureum* | Rhodophyta, Porphyridiales |
| Ct1658 | Chaten1\|1658 | *Chaetoceros tenuissimus* | SAR, Chaetocerotales |
| Bn1393 | Bigna1\|91393 | *Bigelowiella natans* | SAR, Chlorarachniales |
| Pm1017 | Paumic1\|81017 | *Paulinella micropora* | SAR, Euglyphida |
| Pm6562 | Paumic1\|86562 | *Paulinella micropora* | SAR, Euglyphida |
| Ng8424 | Nangad1894_1\|8424 | *Nannochloropsis gaditana* | SAR, Eustigmatales |
| No2883 | Nanoce1779_2\|602883 | *Nannochloropsis oceania* | SAR, Eustigmatales |
| Ae9462 | Alaesc1\|19462 | *Alaria esculenta* | SAR, Laminariales |
| Pt9896 | Phatr2\|9896 | *Phaeodactylum tricornutum* | SAR, Naviculales |
| Ps7279 | Pelago2097_1\|507279 | *Pelagophyceae* sp. | SAR, Pelagomonadales |
| Cc0336 | Cyccr2_2\|10336 | *Cyclotella cryptica* | SAR, Thalassiosirales |
| To8806 | Thaoce1\|88806 | *Thalassiosira oceanica* | SAR, Thalassiosirales |
| Tp2283 | Thaps3\|262283 | *Thalassiosira pseudonana* | SAR, Thalassiosirales |
| Fc8601 | Fracy1\|278601 | *Fragilariopsis cylindrus* | Ochrophyta, Bacillariales |
| Fc8805 | Fracy1\|198805 | *Fragilariopsis cylindrus* | Ochrophyta, Bacillariales |
| Np7520 | Nitput1\|7520 | *Nitzschia putrida* | Ochrophyta, Bacillariales |
| Pi3085 | Parimp1_4\|793085 | *Paraphysomonas imperforata* | Ochrophyta, Chromulinales |
| Pi4313 | Parimp1_4\|664313 | *Paraphysomonas imperforata* | Ochrophyta, Chromulinales |
| Pi6624 | Parimp1_4\|766624 | *Paraphysomonas imperforata* | Ochrophyta, Chromulinales |
| Ms8530 | MonC73_1\|8530 | *Mondopsis strain* | Ochrophyta, Eustigmatales |
| Vs6633 | VisC74_1\|6633 | *Vischeria strain* | Ochrophyta, Eustigmatales |
| Fs1980 | Fisso2\|41980 | *Fistulifera solaris* | Ochrophyta, Naviculales |
| Mp338 | Maypse1\|338 | *Mayamaea pseudoterrestris* | Ochrophyta, Naviculales |
| Sr5136 | Semro1_1\|15136 | *Seminavis robusta* | Ochrophyta, Naviculales |
| Mv1110 | Mintr2\|481110 | *Minidiscus variabilis* | Ochrophyta, Thalassiosirales |
| Ps3250 | Protsul1175_1\|2093250 | *Proteomonas sulcata* | Cryptophyta, Pyrenomonadales |
| Pa2860 | Phaant1\|12860 | *Phaeocystis antarctica* | Haptophyta, Phaeocystales |
| Pa9574 | Phaant1\|39574 | *Phaeocystis antarctica* | Haptophyta, Phaeocystales |
| Oq2912 | Ostque1\|2912 | *Ostreobium quekettii* | Chlorophyta, Bryopsidales |
| Oq8445 | Ostque1\|8445 | *Ostreobium quekettii* | Chlorophyta, Bryopsidales |
| Ts1600 | Tetstr1\|451600 | *Tetraselmis striata* | Chlorophyta, Chlorodendrales |
| Ts1897 | Tetstr1\|451897 | *Tetraselmis striata* | Chlorophyta, Chlorodendrales |
| Ts3708 | Tetstr1\|433708 | *Tetraselmis striata* | Chlorophyta, Chlorodendrales |
| Ts4758 | Tetstr1\|464758 | *Tetraselmis striata* | Chlorophyta, Chlorodendrales |
| Ts8757 | Tetstr1\|428757 | *Tetraselmis striata* | Chlorophyta, Chlorodendrales |
| Ap2048 | Auxeprot1\|2048 | *Auxenochlorella protothecoides* | Chlorophyta, Chlorellales |
| Cs4678 | Chloso1230_1_1\|4678 | *Chlorella sorokiniana* | Chlorophyta, Chlorellales |
| Cv1648 | ChlNC64A_1\|21648 | *Chlorella variabilis* | Chlorophyta, Chlorellales |
| Cv6610 | ChlNC64A_1\|26610 | *Chlorella variabilis* | Chlorophyta, Chlorellales |
| Cv8981 | ChlNC64A_1\|48981 | *Chlorella variabilis* | Chlorophyta, Chlorellales |
| Hs5561 | Heli50920_1\|5561 | *Helicosporidium* sp. | Chlorophyta, Chlorellales |
| Mc1024 | Micco1_1\|1024 | *Micractinium conductrix* | Chlorophyta, Chlorellales |
| Nd7985 | Nandes2437_1\|7985 | *Nannochloris desiccate* | Chlorophyta, Chlorellales |
| Ps4171 | Picsp_1\|4171 | *Picochlorum soloecismus* | Chlorophyta, Chlorellales |
| Ag287 | Astgub1\|287 | *Astrephomene gubernaculifera* | Chlorophyta, Chlamydomonadales |
| Cr1332 | Chlrei\|1332 | *Chlamydomonas reinhardtii* | Chlorophyta, Chlamydomonadales |
| Gp5189 | Gonpec1\|15189 | *Gonium pectoral* | Chlorophyta, Chlamydomonadales |
| Va6426 | Volafr1\|6426 | *Volvox africanus* | Chlorophyta, Chlamydomonadales |
| Vc1055 | Volca2_1\|1055 | *Volvox carteri* | Chlorophyta, Chlamydomonadales |
| Mc281 | MicpuN3v2\|281 | *Micromonas commode* | Chlorophyta, Mamiellales |
| Cz4334 | Chrzof1\|14334 | *Chromochloris zofingiensis* | Chlorophyta, Sphaeropleales |
| Da2672 | DesarB2533_2\|122672 | *Desmodesmus armatus* | Chlorophyta, Sphaeropleales |
| Ec0714 | Enacos1_1\|6790714 | *Enallax costatus* | Chlorophyta, Sphaeropleales |

| | | | |
|---|---|---|---|
| Fr190 | Flerot1_1\|11525190 | *Flechtneria rotunda* | Chlorophyta, Sphaeropleales |
| Mm5487 | Monmin1\|525487 | *Monoraphidium minutum* | Chlorophyta, Sphaeropleales |
| Rs3250 | Rapsub1_1\|13250 | *Raphidocelis subcapitata* | Chlorophyta, Sphaeropleales |
| So3229 | Sceobl\|763229 | *Scenedesmus obliquus* | Chlorophyta, Sphaeropleales |
| Td8890 | Tetrdes\|7768890 | *Tetradesmus deserticola* | Chlorophyta, Sphaeropleales |
| To0874 | Tetrobl72_1\|3960874 | *Tetradesmus obliquus* | Chlorophyta, Sphaeropleales |
| Bb0800 | Botrbrau1_1\|20800 | *Botryococcus braunii* | Chlorophyta, Trebouxiales |
| Cb4441 | Chabra1\|344441 | *Chaura braunii* | Charophyta, Charales |
| Ca1 | Chlat1_1\|1000 | *Chlorokybus atmophyticus* | Charophyta, Chlorokybales |
| Ca1731 | Chrsp_1\|171731 | *Chrysochromulina tobin* | Charophyta, Chlorokybales |
| Ca2577 | Chlat1_1\|2577 | *Chlorokybus atmophyticus* | Charophyta, Chlorokybales |
| Ca8086 | Chlat1_1\|8086 | *Chlorokybus atmophyticus* | Charophyta, Chlorokybales |
| Kn3516 | Klenit1_1\|3516 | *Klebsomidium nitens* | Charophyta, Klebsormidiales |
| Kn4180 | Klenit1_1\|14180 | *Klebsomidium nitens* | Charophyta, Klebsormidiales |
| Mv1 | Mesvir1_1\|15685 | *Mesostigma viride* | Charophyta, Mesostigmatales |
| Sm3175 | Spimu1_1\|23175 | *Spirogloea muscicola* | Charophyta, Spirogloeales |
| Sm4059 | Spimu1_1\|4059 | *Spirogloea muscicola* | Charophyta, Spirogloeales |
| Mk1356 | Meskra\|2261356 | *Mesotaenium kramstae* | Charophyta, Zygnematales |
| Mk5285 | Meskra\|2665285 | *Mesotaenium kramstae* | Charophyta, Zygnematales |
| Zc6981b | Zygcir6981b_2 | *Zygnema circumcarinatum* | Charophyta, Zygnematales |

**Embryophyte ACOSs**

| | | | |
|---|---|---|---|
| Aa213 | Aagr0000081.213 | *Anthoceros agrestis* | Anthocerotophyta, Anthocerotales |
| Mpo0059 | Mapoly0014s0059 | *Marchantia polymorpha* | Marchantiophyta, Marchantiales |
| Cp0400 | CepurGG1.1G280400 | *Ceratodon purpureus* | Bryophyta, Dicranales |
| PpACOS6 | Phpat.003G149600 | *Physcomitrium patens* | Bryophyta, Funariales |
| Pp730 | Pp3c8_730 | *Physcomitrium patens* | Bryophyta, Funariales |
| Sf0036 | Sphfalx0179s0036 | *Sphagnum fallax* | Bryophyta, Sphagnales |
| Tl2900 | Tle3c02900 | *Takakia lepidozioides* | Bryophyta, Takakiales |
| Sm5483 | Sm115483 | *Selaginella moellendorffii* | Lycopodiophyta, Selaginellales |
| Cri41000 | Ceric.33G041000 | *Ceratopteris richardii* | Polypodiopsida, Polypodiales |
| Af3677 | Azfi0027.g023677 | *Azolla filiculoides* | Polypodiopsida, Salviniales |
| Sc1820 | Sacu0127.g021820 | *Salvinia cucullate* | Polypodiopsida, Salviniales |
| GbACOS | JANKJI010000003 | *Gingko biloba* | Ginkgophyta, Ginkgoales |
| CjACOS | BDR24742 | *Cryptomeria japonica* | Gymnosperms, Cupressales |
| Tp0007 | Thupl.29382414s0007 | *Thuja plicata* | Gymnosperms, Cupressales |
| Gmo2049 | MNCI01032049 | *Gnetum monatum* | Gymnosperms, Gnetales |
| Atr297 | Atscaffold00025.297 | *Amborella trichopoda* | Basal angiosperms, Amborellales |
| Nc2160 | Nycol.B02160 | *Nymphaea colorata* | Basal angiosperms, Nymphaeales |
| Ck0900 | CKAN_01300900 | *Cinnamomum kanehirae* | Core angiosperms, Laurales |
| Cs0014 | CsJAJIZI010000014 | *Chloranthus sessilifolius* | Core angiosperms, Chloranthales |
| Bv3515 | BvEL10Ac6g13515 | *Beta vulgaris* | eudicots, Caryophyllales |
| Ls8221 | Ls148221 | *Lactuca sativa* | eudicots, asterids, Asterales |
| St1030 | Soltu.DM.02G031030.2 | *Solanum tuberosum* | eudicots, asterids, Solanales |
| Ns3489 | AY163489 | *Nicotiana sylvestris* | eudicots, asterids, Solanales |
| AtACOS5 | At1G62940 | *Arabidopsis thaliana* | eudicots, rosids, Brassicales |
| BnACOS5 | XP_013664377 | *Brassica napus* | eudicots, rosids, Brassicales |
| Ah7DH | AhWWT7DH | *Arachis hypogaea* | eudicots, rosids, Fabales |
| Gm9700 | Glyma.08G329700 | *Glycine max* | eudicots, rosids, Fabales |
| Gm6900 | Glyma.18G076900 | *Glycine max* | eudicots, rosids, Fabales |
| Mt0770 | Medtr3g460770 | *Medicago truncatula* | eudicots, rosids, Fabales |
| PtACOS13 | Potri.001G055700 | *Populus trichocarpa* | eudicots, rosids, Malpighiales |
| Md0400 | MD13G1120400 | *Malus domestica* | eudicots, rosids, Rosales |
| Aa2200 | Acora.06G002200 | *Acorus americanus* | Basal monocots, Acorales |
| Sp4500 | Spipo12G0064500 | *Spirodela polyrhiza* | Basal monocots, Alismatales |
| OsACOS12 | Os04g24530 | *Oryza sativa* | monocots, Poales |
| Zm3702 | Zm00001d003702_P001 | *Zea mays* | monocots, Poales |
| Cn4391 | CnWOK94391 | *Canna indica* | monocots, Zingiberales |
| Ma6020 | Ma6P36020_001 | *Musa acuminata* | monocots, Zingiberales |

**Embryophyte 4CLs**

| | | | |
|---|---|---|---|
| Aa18 | AagrOXF000004l.18 | *Anthoceros agrestis* | Anthocerotophyta, Anthocerotales |
| Mp0014 | Mapoly0197s0014 | *Marchantia polymorpha* | Marchantiophyta, Marchantiales |
| Pp4CL1 | Phpat.021G054800 | *Physcomitrium patens* | Bryophyta, Funariales |
| Pp4CL2 | Phpat.018G019800 | *Physcomitrium patens* | Bryophyta, Funariales |
| Pp4CL3 | Phypa1_1\|104959 | *Physcomitrium patens* | Bryophyta, Funariales |
| Pp4CL4 | Phpat.022G056000 | *Physcomitrium patens* | Bryophyta, Funariales |
| Tl3606 | Tle1c03606 | *Takakia lepidozioides* | Bryophyta, Takakiales |
| Tl7268 | Tle1c07268 | *Takakia lepidozioides* | Bryophyta, Takakiales |
| Sm1251 | Sm171251 | *Selaginella moellendorffii* | Lycopodiophyta, Selaginellales |
| Sm3133 | Sm173133 | *Selaginella moellendorffii* | Lycopodiophyta, Selaginellales |
| Sm7393 | Sm177393 | *Selaginella moellendorffii* | Lycopodiophyta, Selaginellales |
| Cri3800 | Ceric.17G083800 | *Ceratopteris richardii* | Polypodiopsida, Polypodiales |
| Cri5100 | Ceric.1Z155100 | *Ceratopteris richardii* | Polypodiopsida, Polypodiales |
| Cri8400 | Ceric.15G048400 | *Ceratopteris richardii* | Polypodiopsida, Polypodiales |
| Cri8600 | Ceric.12G038600 | *Ceratopteris richardii* | Polypodiopsida, Polypodiales |
| Tp0001 | Thupl29343480s0001 | *Thuja plicata* | Gymnosperms, Cupressales |
| Tp0002 | Thupl29377326s0002 | *Thuja plicata* | Gymnosperms, Cupressales |
| Tp0003 | Thupl29351263s0003 | *Thuja plicata* | Gymnosperms, Cupressales |
| Tp0008 | Thupl29379587s0008 | *Thuja plicata* | Gymnosperms, Cupressales |
| Tp0029 | Thupl29378212s0029 | *Thuja plicata* | Gymnosperms, Cupressales |
| Atr261 | Atscaffold00019.261 | *Amborella trichopoda* | Basal angiosperms, Amborellales |
| Atr51 | Atscaffold00048.51 | *Amborella trichopoda* | Basal angiosperms, Amborellales |
| Msi0009 | MsJARNKS010000009 | *Magnolia sinica* | Core angiosperms, Magnoliales |
| Msi3776 | MsXP_058113776 | *Magnolia sinica* | Core angiosperms, Magnoliales |
| Msi4730 | MsXP_058084730 | *Magnolia sinica* | Core angiosperms, Magnoliales |
| Msi7945 | MsXP_058067945 | *Magnolia sinica* | Core angiosperms, Magnoliales |
| At4CL1 | AT1G51680 | *Arabidopsis thaliana* | eudicots, rosids, Brassicales |
| At4CL2 | AT3G21240 | *Arabidopsis thaliana* | eudicots, rosids, Brassicales |
| At4CL3 | AT1G65060 | *Arabidopsis thaliana* | eudicots, rosids, Brassicales |
| At4CL4 | AT3G21230 | *Arabidopsis thaliana* | eudicots, rosids, Brassicales |
| Pt4CL2 | Potri.018G094200 | *Populus trichocarpa* | eudicots, rosids, Malpighiales |
| Pt4CL3 | Potri.019G049500 | *Populus trichocarpa* | eudicots, rosids, Malpighiales |
| Pt4CL4 | Potri.003G188500 | *Populus trichocarpa* | eudicots, rosids, Malpighiales |
| Pt4CL5 | Potri.001G036900 | *Populus trichocarpa* | eudicots, rosids, Malpighiales |
| Sp0700 | Spipo10G0010700 | *Spirodela polyrhiza* | Basal monocots, Alismatales |
| Sp3800 | Spipo12G0053800 | *Spirodela polyrhiza* | Basal monocots, Alismatales |
| Sp7700 | Spipo5G0067700 | *Spirodela polyrhiza* | Basal monocots, Alismatales |
| Zma0220 | Zosma182g00220 | *Zostera marina* | Basal monocots, Alismatales |
| Zma0620 | Zosma10g00620 | *Zostera marina* | Basal monocots, Alismatales |
| Zma1830 | Zosma7g01830 | *Zostera marina* | Basal monocots, Alismatales |
| Os4CL1 | Os08g14760 | *Oryza sativa* | monocots, Poales |
| Os4CL2 | Os02g46970 | *Oryza sativa* | monocots, Poales |
| Os4CL3 | Os02g08100 | *Oryza sativa* | monocots, Poales |
| Os4CL4 | Os06g44620 | *Oryza sativa* | monocots, Poales |
| Os4CL5 | Os08g34790 | *Oryza sativa* | monocots, Poales |
| Zo0987 | ZXP_042410987 | *Zingiber officinale* | monocots, Zingiberales |
| Zo3702 | ZXP_042393702 | *Zingiber officinale* | monocots, Zingiberales |
| Zo4244 | ZXP_042444244 | *Zingiber officinale* | monocots, Zingiberales |
| Zo5748 | ZXP_042415748 | *Zingiber officinale* | monocots, Zingiberales |

### Table S2 Sequences used for CCR family tree construction (Fig S2)

| Name | Gene ID | Species | Classification |
|---|---|---|---|
| **Algal homologues** | | | |
| Bn8692 | Bigna1\|78692 | *Bigelowiella natans* | SAR, Chlorarachniales |
| Cv1077 | Chrveli1\|1077 | *Chromera velia* | SAR, Chromerida |
| Pm5788 | Paumic1\|65788 | *Paulinella micropora* | SAR, Euglyphida |
| No4910 | Nanoce84910 | *Nannocholoropsis oceania* | SAR, Eustigmatales |
| Fc8633 | Fracro1\|8633 | *Fragilaria crotonensis* | SAR, Fragilariales |
| Mp6776 | Macpyr2\|9466776 | *Macrocystis pyrifera* | SAR, Laminariales |
| Pt1257 | Phatr2\|31257 | *Phaeodactylum tricornutum* | SAR, Naviculales |
| Ms7850 | MonC141\|7850 | *Mondopsis strain* | Ochrophyta, Eustigmatales |
| Ms9869 | MonC73\|9869 | *Mondopsis strain* | Ochrophyta, Eustigmatales |
| Vs7887 | VisC74\|7887 | *Vischeria strain* | Ochrophyta, Eustigmatales |
| Cs2293 | Crypto2293 | *Cryptophyceae* sp. | Cryptista, Pyrenomonadales |
| Gt1113 | Guith1\|1113 | *Guilardia theta* | Cryptophyta, Pyrenomonadales |
| Ps1175 | Protsul11755 | *Proteomonas sulcata* | Cryptophyta, Pyrenomonadales |
| Cp9666 | Chrpa1\|19666 | *Chrysochromulina parva* | Haptophyta, Prymnesiales |
| Oq4852 | Ostque1\|4852 | *Ostreobium quekettii* | Chlorophyta, Bryopsidales |
| Cr4532 | ChlreiCC4532 | *Chlamydomonas reinhardtii* | Chlorophyta, Chlamydomonadales |
| Gp1354 | Gonpec1\|1354 | *Gonium pectoral* | Chlorophyta, Chlamydomonadales |
| Nd576 | Nandes2437\|5 | *Nannochloris desiccata* | Chlorophyta, Chlorellales |
| Cz6754 | Chrzof1\|6754 | *Chromochloris zofingiensis* | Chlorophyta, Sphaeropleales |
| Da2533 | DesarB2533 | *Desmodesmus armatus* | Chlorophyta, Sphaeropleales |
| Es3294 | Enacos1\|6453294 | *Enallax costatus* | Chlorophyta, Sphaeropleales |
| Fr2494 | Flerot1\|10932494 | *Flechtneria rotunda* | Chlorophyta, Sphaeropleales |
| Mm4590 | Monmin1\|384590 | *Monoraphidium minutum* | Chlorophyta, Sphaeropleales |
| Bb3222 | Botrbrau1\|13222 | *Botryococcus braunii* | Chlorophyta, Trebouxiales |
| Ts4402 | Trebou4402 | *Trebouxiophyceae* sp. | Chlorophyta, Trebouxiales |
| Ca6773 | Chlat1\|6773 | *Chlorokybus atmophyticus* | Charophyta, Chlorokybales |
| Kn3226 | Klenit1\|13226 | *Klebsomidium nitens* | Charophyta, Klebsormidiales |
| Zc6981 | Zygcir6981 | *Zygnema circumcarinatum* | Charophyta, Zygnematales |
| Mk657 | Meskra657 | *Mesotaenium kramstae* | Charophyta, Zygnematales |
| **Embryophytes TKPRs** | | | |
| Aa144 | Aagr0000381.144 | *Anthoceros agrestis* | Anthocerotophyta, Anthocerotales |
| Aa275 | Aagr0000631.275 | *Anthoceros agrestis* | Anthocerotophyta, Anthocerotales |
| Mpo0037 | Mapoly0028s0037 | *Marchantia polymorpha* | Marchantiophyta, Marchantiales |
| Mpo0025 | Mapoly0077s0025 | *Marchantia polymorpha* | Marchantiophyta, Marchantiales |
| Cp3100 | Cepur3G083100 | *Ceratodon purpureus* | Bryophyta, Dicranales |
| Cp9600 | Cepur12G099600 | *Ceratodon purpureus* | Bryophyta, Dicranales |
| PpTKPR1 | Pp3c16_1620 | *Physcomitrium patens* | Bryophyta, Funariales |
| PpTKPR2 | Pp3c8_16620 | *Physcomitrium patens* | Bryophyta, Funariales |
| Sf0028 | Sphfalx0044s0028 | *Sphagnum fallax* | Bryophyta, Sphagnales |
| Sf0290 | Sphfalx0000s0290 | *Sphagnum fallax* | Bryophyta, Sphagnales |
| Tl6940 | Tle1c06940 | *Takakia lepidozioides* | Bryophyta, Takakiales |
| Tl1087 | Tle1c001087 | *Takakia lepidozioides* | Bryophyta, Takakiales |
| Sm74610 | Sm\|74610 | *Selaginella moellendorffii* | Lycopodiophyta, Selaginellales |
| Sm80798 | Sm\|80798 | *Selaginella moellendorffii* | Lycopodiophyta, Selaginellales |
| Cri2900 | Ceric.32G062900 | *Ceratopteris richardii* | Polypodiopsida, Polypodiales |
| Cri73400 | Ceric.32G073400 | *Ceratopteris richardii* | Polypodiopsida, Polypodiales |
| Af3772 | Azfi043772 | *Azolla filiculoides* | Polypodiophyta, Salviniales |
| Af0318 | Azfi060318 | *Azolla filiculoides* | Polypodiophyta, Salviniales |
| Sc3701 | Sacu023701 | *Salvinia cucullata* | Polypodiophyta, Salviniales |
| Sc7512 | Sacu007512 | *Salvinia cucullata* | Polypodiophyta, Salviniales |
| Gb31436 | Gb_31436 | *Ginkgo biloba* | Ginkgophyta, Ginkgoales |
| Tp0003 | Thupl29378135s0003 | *Thuja plicata* | Gymnosperms, Cupressales |
| GmoTKPR | MNCI01015527 | *Gnetum monatum* | Gymnosperms, Gnetales |

| | | | |
|---|---|---|---|
| Atr248 | Atri00005.248 | *Amborella trichopoda* | Basal angiosperms, Amborellales |
| Atr35 | Atri00022.35 | *Amborella trichopoda* | Basal angiosperms, Amborellales |
| Nc0883 | NycolF00883 | *Nymphaea colorata* | Basal angiosperms, Nymphaeales |
| Nc1577 | NycolA01577 | *Nymphaea colorata* | Basal angiosperms, Nymphaeales |
| Ck2300 | CKAN01442300 | *Cinnamomum kanehirae* | Core angiosperms, Laurales |
| Ck2700 | CKAN00102700 | *Cinnamomum kanehirae* | Core angiosperms, Laurales |
| MsiTKPR2 | MsTKPRR2 | *Magnolia sinica* | Core angiosperms, Magnoliales |
| MsiTKPR1 | MsTKPR1 | *Magnolia sinica* | Core angiosperms, Magnoliales |
| Cs0001 | CsJAJIZI010000001 | *Chloranthus sessilifolius* | Core angiosperms, Chloranthales |
| Bv4416 | Bv6g14416 | *Beta vulgaris* | eudicots, Caryophyllales |
| Bv1573 | Bv9g21573 | *Beta vulgaris* | eudicots, Caryophyllales |
| Ls59300 | Ls59300 | *Lactuca sativa* | eudicots, asterids, Asterales |
| Ls48581 | Ls48581 | *Lactuca sativa* | eudicots, asterids, Asterales |
| St3904 | StDMP400043904 | *Solanum tuberosum* | eudicots, asterids, Solanales |
| Gm7200 | Glyma07G157200 | *Glycine max* | eudicots, rosids, Fabales |
| Gm5600 | Glyma13G355600 | *Glycine max* | eudicots, rosids, Fabales |
| AhB9NP0R | AhB9NP0R | *Arachis hypogaea* | eudicots, rosids, Fabales |
| AhDVK8Y6 | AhDVK8Y6 | *Arachis hypogaea* | eudicots, rosids, Fabales |
| Md2500 | Md13G1042500 | *Malus domestica* | eudicots, rosids,Rosales |
| Md5200 | Md10G1145200 | *Malus domestica* | eudicots, rosids,Rosales |
| AtTKPR2 | AT1G68540 | *Arabidopsis thaliana* | eudicots, rosids, Brassicales |
| AtTKPR1 | AT4G35420 | *Arabidopsis thaliana* | eudicots, rosids, Brassicales |
| Ac1600 | Acora11G091600 | *Acorus americanus* | Basal monocots, Acorales |
| Ac0500 | Acora09G150500 | *Acorus americanus* | Basal monocots, Acorales |
| Sp6700 | Spipo10G0016700 | *Spirodela polyrhiza* | Basal monocots, Alismatales |
| Os3670 | Os01g03670 | *Oryza sativa* | monocots, Poales |
| Os0440 | Os08g40440 | *Oryza sativa* | monocots, Poales |
| Zm0970 | Zmd020970_P001 | *Zea mays* | monocots, Poales |
| Zm1488 | Zmd031488_P001 | *Zea mays* | monocots, Poales |
| CnTKPR | CnTKPR | *Canna indica* | monocots, Zingiberales |
| Ma6P0 | Ma6P09630_001 | *Musa acuminata* | monocots, Zingiberales |
| MaP11 | MaP11930_001 | *Musa acuminata* | monocots, Zingiberales |
| **CCRL** | | | |
| Aa87 | Aagr0000241.87 | *Anthoceros agrestis* | Anthocerotophyta, Anthocerotales |
| Pp2220 | Pp3c7_12220 | *Physcomitrium patens* | Bryophyta, Funariales |
| Tl2164 | Tle1c02164 | *Takakia lepidozioide* | Bryophyta, Takakiales |
| AtCCRL14 | AT5G19440 | *Arabidopsis thaliana* | eudicots, rosids, Brassicales |
| Os4480 | Os01g34480 | *Oryza sativa* | monocots, Poales |
| **CCR** | | | |
| Mpo0049 | Mapoly0063s0049 | *Marchantia polymorpha* | Marchantiophyta, Marchantiales |
| Pp1820 | Pp3c1_1820 | *Physcomitrium patens* | Bryophyta, Funariales |
| Pp2950 | Pp3c11_2950 | *Physcomitrium patens* | Bryophyta, Funariales |
| Pp4520 | Pp3c2_34520 | *Physcomitrium patens* | Bryophyta, Funariales |
| PpCCR | Pp3c7_17190 | *Physcomitrium patens* | Bryophyta, Funariales |
| Tl643 | Tle2c02643 | *Takakia lepidozioides* | Bryophyta, Takakiales |
| Tl6214 | Tle1c06214 | *Takakia lepidozioides* | Bryophyta, Takakiales |
| Sm141996 | Sm141996 | *Selaginella moellendorffii* | Lycopodiophyta, Selaginellales |
| Sm271114 | Sm271114 | *Selaginella moellendirffii* | Lycopodiophyta, Selaginellales |
| Cri4000 | Ceric.13G084000 | *Ceratopteris richardii* | Polypodiopsida, Polypodiales |
| Cri3400 | Ceric.08G023400 | *Ceratopteris richardii* | Polypodiopsida, Polypodiales |
| Tp0008 | Thupl29380998s0008 | *Thuja plicata* | Gymnosperms, Cupressales |
| Atr159 | Atri00065.159 | *Amborella trichopoda* | Basal angiosperms, Amborellales |
| Msi7625 | Ms058067625 | *Magnolia sinica* | Core angiosperms, Magnoliales |
| AtCCR1 | AT1G15950 | *Arabidopsis thaliana* | eudicots, rosids, Brassicales |
| PtCCR2 | PtCCR2 | *Populus trichocarpa* | eudicots, rosids, Malpighiales |
| Sp5100 | Spipo0G0185100 | *Spirodela polyrhiza* | monocots, Alismatales |
| Zma350 | Zosma16g01350 | *Zostera marina* | monocots, Alismatales |
| Os4050 | Os09g04050 | *Oryza sativa* | monocots, Poales |
| Os4280 | Os08g34280 | *Oryza sativa* | monocots, Poales |

| | | | |
|---|---|---|---|
| Os5150 | Os09g25150 | *Oryza sativa* | monocots, Poales |
| Os8420 | Os02g08420 | *Oryza sativa* | monocots, Poales |
| Zo4428 | Z0042464428 | *Zingiber officinale* | monocots, Zingiberales |
| **ANR** | | | |
| Atr30 | Atri00098.30 | *Amborella trichopoda* | Basal angiosperms, Amborellales |
| Msi2931 | Ms058072931 | *Magnolia sinica* | Core angiosperms, Magnoliales |
| AtANR | AT1G61720 | *Arabidopsis thaliana* | eudicots, rosids, Brassicales |
| PtANR1 | PtANR1 | *Populus trichocarpa* | eudicots, rosids, Malpighiales |
| Sp4900 | Spipo14G0054900 | *Spirodela polyrhiza* | monocots, Alismatales |
| Os3800 | Os04g53800 | *Oryza sativa* | monocots, Poales |
| Os3810 | Os04g53810 | *Oryza sativa* | monocots, Poales |
| Os3850 | Os04g53850 | *Oryza sativa* | monocots, Poales |
| Os3920 | Os04g53920 | *Oryza sativa* | monocots, Poales |
| Zo6220 | Z042386220 | *Zingiber officinale* | monocots, Zingiberales |
| **DFR** | | | |
| Tp0006 | Thupl29378220s0006 | *Thuja plicata* | Gymnosperms, Cupressales |
| Tp0015 | Thupl29378454s0015 | *Thuja plicata* | Gymnosperms, Cupressales |
| Atr95 | Atri00011.95 | *Amborella trichopoda* | Basal angiosperms, Amborellales |
| Msi8380 | Ms058088380 | *Magnolia sinica* | Core angiosperms, Magnoliales |
| AtDFR | AT5G42800 | *Arabidopsis thaliana* | eudicots, rosids, Brassicales |
| At7250 | AT4G27250 | *Arabidopsis thaliana* | eudicots, rosids, Brassicales |
| PtDFR | PtDFR | *Populus trichocarpa* | eudicots, rosids, Malpighiales |
| Sp0200 | Spipo10G0000200 | *Spirodela polyrhiza* | monocots, Alismatales |
| Zma730 | Zosma1g00730 | *Zostera marina* | monocots, Alismatales |
| Os4260 | Os01g44260 | *Oryza sativa* | monocots, Poales |
| Zo8899 | Z042378899 | *Zingiber officinale* | monocots, Zingiberales |

**Fig. S1 Maximum Likelihood tree of embryophyte ACOS and 4CL sequences and algal homologues**

Included in the tree are ACOSs and 4CLs from representative embryophytes from all major lineages from hornworts to eudicots and 76 algal homologues that scored BLASTp E-values smaller than $1{\times}e^{-20}$ (Table S1). An unrooted ML tree was constructed using MEGA ver. 11 with the WAG+G+I substitution model. Support for the inferred tree was estimated using 1000 bootstrap replicates and bootstrap values (>50%) are displayed at the nodes.

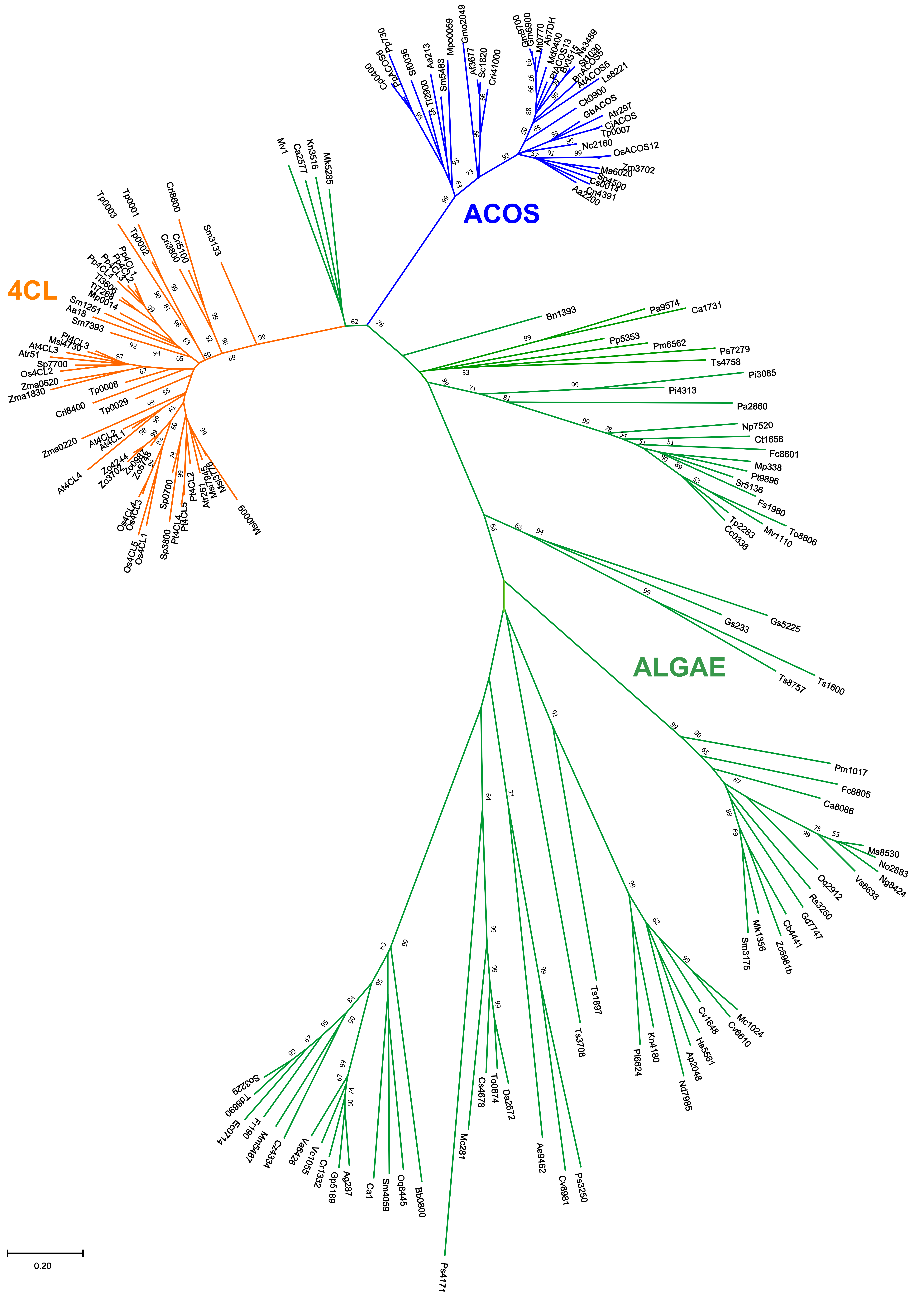
ACOS
4CL
ALGAE
0.20
Cp0400
PpACOS6
Pp730
Sf0036
Tl2900
Aa213
Sm5483
Mpo0059
Gmo2049
At3677
Sc1820
Cri41000
Gm9700
Gm6900
Mt0770
Ah7DH
Md0400
PtACOS13
Bv3515
Ns3489
Sl1030
BnACOS5
AtACOS5
Ls8221
Ck0900
GbACOS
Atr297
CjACOS
Tp0007
Nc2160
OsACOS12
Zm3702
Ma6020
Sp4500
Cs0014
Cn4391
Aa2200
Mv1
Ca2577
Kn3516
Mk5285
Tp0003
Tp0001
Tp0002
Cri8600
Cri5100
Cri3800
Sm3133
Pp4CL1
Pp4CL2
Pp4CL3
Pp4CL4
Tl3606
Tl7268
Mp0014
Sm1251
Aa18
Sm7393
Pt4CL3
Msi4730
At4CL3
Atr51
Sp7700
Os4CL2
Zma0620
Zma1830
Tp0008
Cri8400
Tp0029
Zma0220
At4CL2
At4CL1
At4CL4
Zo4244
Zo3702
Zo0987
Zo5746
Os4CL4
Os4CL3
Os4CL5
Os4CL1
Sp0700
Sp3800
Pt4CL4
Pt4CL5
Pt4CL2
Atr261
Msi7945
Msi5776
Msi0009
Bn1393
Pa9574
Ca1731
Pp5353
Pm6562
Ps7279
Ts4758
Pi3085
Pi4313
Pa2860
Np7520
Ct1658
Fc8601
Mp338
Pt9896
Sr5136
Fs1980
Tp2283
Cc0336
Mv1110
To8806
Gs233
Gs5225
Ts8757
Ts1600
Pm1017
Fc8805
Ca8086
Ms8530
No2883
Ng8424
Vs6633
Oq2912
Rs3250
Gd7747
Cb4441
Zc6981b
Mk1356
Sm3175
Ts1897
Ts3708
Kn4180
Pl6624
Nd7985
Ap2048
Hs5561
Cv1648
Cv6610
Mc1024
Cs4678
To0874
Da2672
Ae9462
Cv8681
Ps3250
Mc281
Ps4171
So3229
Td8890
Ec0714
Ff190
Mm5487
Cz4334
Va6426
Vc1055
Cr1332
Gp5189
Ag287
Ca1
Sm4059
Oq8445
Bb0800

**Fig. S2 Maximum Likelihood tree of TKPR and other enzymes of the CCR family and algal homologues**

Included in the tree are TKPRs from thirty-five embryophytes representing all major lineages from hornworts to eudicots, CCR, CCRL, DFR and ANR from select embryophytes, and 29 algal homologues that scored BLASTp E-values smaller than $1 \times e^{-20}$ (Table S2). An unrooted ML tree was constructed using MEGA ver. 11 with the WAG+G+I substitution model. Support for the inferred tree was estimated using 1000 bootstrap replicates and bootstrap values (>50%) are displayed at the nodes.

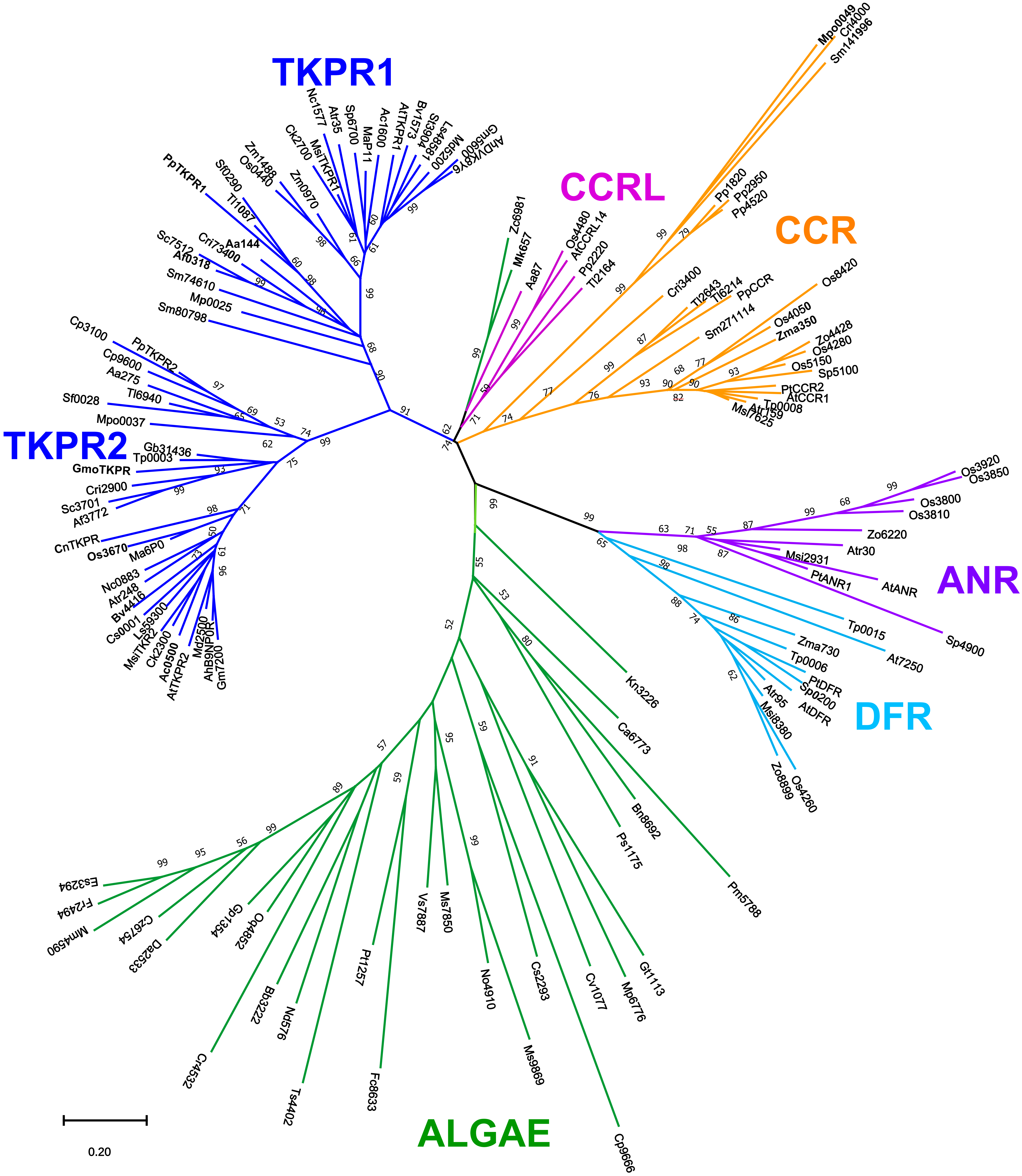

TKPR1
CCRL
CCR
TKPR2
ANR
DFR
ALGAE
0.20
Mpo0049
Cri4000
Sm141996
Pp1820
Pp2950
Pp4520
Cri3400
Tl2643
Tl6214
PpCCR
Sm271114
Os8420
Os4050
Zma350
Zo4428
Os4280
Os5150
Sp5100
PtCCR2
AtCCR1
Tp0008
Atr159
Msi7625
Zc6981
Mk657
Aa87
Os4480
AtCCRL14
Pp2220
Tl2164
Nc1577
Atr35
Sp6700
MaP11
Ac1600
AtTKPR1
Bv1573
Sl3904
Ls4858
Md5200
Gm5600
AhDVK8Y6
MsiTKPR1
Ck2700
Zm0970
Zm1488
Os0440
Sf0290
Tl1087
PpTKPR1
Aa144
Cri73400
Sc7512
Af0318
Sm74610
Mp0025
Sm80798
Cp3100
PpTKPR2
Cp9600
Aa275
Tl6940
Sf0028
Mpo0037
Gb31436
Tp0003
GmoTKPR
Cri2900
Sc3701
Af3772
CnTKPR
Os3670
Ma6P0
Nc0883
Atr248
Bv4416
Cs0001
Ls59300
MsiTKR2
Ck2300
Ac0500
AtTKPR2
Md2500
AhB9NP0R
Gm7200
Os3920
Os3850
Os3800
Os3810
Zo6220
Atr30
Msi2931
PtANR1
AtANR
Sp4900
Tp0015
At7250
Zma730
Tp0006
PtDFR
Sp0200
AtDFR
Atr95
Msi8380
Zo8899
Os4260
Kn3226
Ca6773
Bn8692
Ps1175
Pm5788
Es3294
Fr2494
Mm4590
Cz6754
Da2533
Gp1354
Oq4852
Cr4532
Bb5222
Nd576
Ts4402
Pt1257
Fc8633
Vs7887
Ms7850
No4910
Ms9869
Cs2293
Cv1077
Mp6776
Gt1113
Cp9666

**Fig. S3 Multiple sequence alignment of ACOSs, 4CLs and algal homologues**

Amino acid residues that form the binding pocket for phenolate substrate (e.g. hydroxycinnamate, ferulate) and determine substrate specificity of 4CL are indicated by diamonds, and those in close contact with the substrate are highlighted in yellow.[1–3] ACOS consensus residues that are predicted to be at the substrate binding pocket and uniquely different from those in 4CL are highlighted in green. Enzymes are from Table S1 and Fig S1 except for Nt4CL2 (*Nicotiana tabacum* 4CL2, accession number AAB18368). Three homologous sequences from Charophyte algae (in green) are also aligned.

```
                    ♦♦♦♦♦♦♦
Nt4CL2       LCVLPLFHIYSLNSVLLCGLRVGAAILIMQKFDIVSFLELIQRYKVTIGPFVPPIVLAIA 289
At4CL4       LCFLPMFHIYALDALMLSAMRTGAALLIVPRFELNLVMELIQRYKVTVVPVAPPVVLAFI 318
Pt4CL4       LCVLPMFHIYALNSIMLCGLRVGAAILIMPKFEIGSLLGLIEKYKVSIAPVVPPVMVAIA 290
Os4CL2       LCVLPLFHIFSLNSVLLCAVRAGAAVALMPRFEMGAMLGAIERWRVTVAAVVPPLVLALA 319
Pp4CL1       MCVLPMFHIYSLNSILLCGLRVGATLVIMPKFELSKMLELIQKHKVTMGPFVPPIVLAIA 323
PpACOS6      LGLMPFFHIYGICGICCAAMRLKGKVVVMARYNFQEFLDILLKYEITFAPIVPPILLQLV 315
SmACOS       LGLMPFFHIYGITGIGCSTMRLKGKVVVMERYELRKMLEALIKYEVQFAPLVPPIILSLV 296
AtACOS5      LGLIPFFHIYGIVGICCATMKNKGKVVAMSRYDLRIFLNALIAHEVSFAPIVPPIILNLV 287
PtACOS13     LGLIPFFHIYGITGICCATLRNKGKVVVMGRFELRTFLNALITQEVSFAPIVPPIILALV 337
OsACOS12     LGLMPFFHIYGITGICCATLRHKGTVVVMDRFDLRTFLRALVDHRVMFAPLVPPVMLAMV 298
Klenit3516   LGLLPFFHIYGMTTLILSPLVKCIPLVVMPKFDLVEMLELIQKYKISFANLVPPVILALA 285
Zygcir6981b  LALLPMFHIYAMTCINMVALKAGATLVTMPRFDLEHMLTLGEKYGITMAPLVPPIILALA 306
Meskra5285   LALLPMFHIYAITAVNMMALKTGATLVTMARFDMEQMLQLAEKYCITIAPVVPPLVLALA 224
```

```
                               ♦♦♦♦                      ♦♦♦♦♦♦♦♦♦♦♦♦♦
Nt4CL2       KSPMVDDYDLS--SVRTVMSGAAPLGKELEDTVRAKFPNAKLGQGYGMTEAGPVLAMCLA 347
At4CL4       KSPETERYDLS--SVRIMLSGAATLKKELEDAVRLKFPNAIFGQGYGMTESG-TVAKSLA 375
Pt4CL4       KSPDLDKHDLS--SLRMLKSGGSPLGKELEDTVRARFPQARLGQGYGMTEAGPVLAMCLA 348
Os4CL2       KNPFVERHDLS--SIRIVLSGAAPLGKELEDALRARLPQAIFGQGYGMTEAGPVLSMCPA 377
Pp4CL1       KNPIVENYDLS--SIKMVMSGAAPLGKELEDAFRARLPNAVLGQGYGMTEAGPVLAMCLA 381
PpACOS6      KKDLGENFDRSKLKLKSILTAAAPLGIELQRAFEAKFPGVEVQQAYGLTEYSCVTVSHCS 375
SmACOS       KNPLVDEYDLSKLKLNAVMTAAAPLAPELQRAFEAKFPGVEMRQAYGLTEYSCVTLSHCA 356
AtACOS5      KNPIVDEFDLSKLKLQSVMTAAAPLAPELLTAFEAKFPNVQVQEAYGLTEHSCITLTHGD 347
PtACOS13     KNPIVEEFDLSKLKLKAIMTAAAPLAPELLTAFENKFPGVQVQEAYGLTEHSCITLTHGD 397
OsACOS12     KSPVADEFDLSDLALKSVMTAAAPLAPDLLAAFQRKFPGVQVEEAYGLTEHSCITLTHAA 358
Klenit3516   KHPVIDKYDLS--SMRGIMSGAAPLGKELQQAAAKRLN-VTVAQGYGMTESSPITLVTRN 342
Zygcir6981b  KSSVVNKYKWH--SLKKVVSGAAPLGADLELAFRSRFPKVVIGQ------SSPTLTLSRG 358
Meskra5285   KSPLVNKYKWS--SLKRVVSGAAPLGAELELVFRKRFPHVAIGQGYGMTESSPTVTLSRA 282
```


[1] Schneider K, Hövel K, Witzel K, Hamberger B, Schomburg D, Kombrink E, Stuible H-P. 2003. The substrate specificity-determining amino acid code of 4-coumarate:CoA ligase. *Proc Natl Acad Sci U S A* 100: 8601–8606.

[2] Hu Y, Gai Y, Yin L, Wang X, Feng C, Feng L, Li D, Jiang X-N, Wang D-C. 2010. Crystal structures of a *Populus tomentosa* 4-coumarate:CoA ligase shed light on its enzymatic mechanisms. *Plant Cell* 22: 3093–3104.

[3] Li Z, Nair SK. 2015. Structural basis for specificity and flexibility in a plant 4-coumarate:CoA ligase. Structure 23: 2032–2042.

**Fig. S4 Multiple sequence alignment of representative enzymes of the embryophyte CCR family and algal homologues**

TKPR sequences are in blue, while DFR (dihydroflavonol 4-reductase), ANR (anthocyanidin reductase), CCR (cinnamoyl-CoA reductase) and CCR-like sequences are in black. Two homologous sequences (in green) from Zygnematales that are closer to embryophyte CCR sequences in a phylogenetic tree (Fig S2) are also aligned. Amino acid residues that form binding pockets for phenolic substrates in DFR,[1] ANR[2] and CCR[3,4] are highlighted in yellow. In particular, Ser/Tyr of ANR and Asp/Asn of DFR (underlined) are critical in determining substrate specificity of these enzymes. The conserved Arg of TKPR (Arg[137] of PpTKPR1) aligns with the Ser and Asp/Asn and therefore is expected to play a critical role in the substrate specificity of TKPR (highlighted in green). Also, Ser[197] (or Thr in some TKPRs) is conserved in TKPR, while the corresponding residue in CCR is nonpolar V or P. Enzymes are listed in Table S2.

```
AtANR        CLKS-KSVKRVIYTSSAAAVSINN---LSGTGIVMNEENWTDVEFLTEEKPFNWGYPISK 173
SbANR        CVKA-GTVRRVILTSSAAGVYIRP--DLQGDGHALDEDSWSDVDFLRANKPPTWGYCVSK 175
PtDFR        CANS-KTVRKIVFTSSAGTVDVEE---KRKP--VYDESCWSDLDFVQSIKMTGWMYFVSK 167
AtDFR        CVKA-KTVRRFVFTSSAGTVNVEE---HQKN--VYDENDWSDLEFIMSKKMTGWMYFVSK 167
PvDFR        CKEA-GTVRRVVFTSTAGAVNVEE---RQKP--VYDENNWSDVDFCRRVKMTGWMYFVSK 182
AtCCRL14     CAKASSVKRVVV-TSSMAAVGYNG--KPRTPDVTVDETWFSDPELCEASK---MWYVLSK 168
MtCCR        SA--EAKVRRVVFTSSIGTVYMDP---NTSRDVVVDESYWSDLEHCKNTK---NWYCYGK 169
AtCCR1       AA--EAKVKRVVITSSIGAVYMDP---NRDPEAVVDESCWSDLDFCKNTK---NWYCYGK 165
PpCCR        CK--EAHVKRVVMTSSIGAVYMNP---SIQPDQEVDESCWSDEAFLRGRK---EWYCLAK 276
AaTKPR2      ------SVKRVVMTSSCSAIRYD--YGHSEEEPPLDESTWSNVEYCKEYK---LWYPLAK 109
GbTKPR       CSRS-PSVKRVVLTSSCSSIRYD--YNNTQHSP-LNETYWSNPEYCKQHN---LWYAYAK 163
Os01g03670   CARASPRPRRVVFTSSCSCVRY----GAGAAAA-LNESHWSDAAYCAAHG---LWYAYAK 176
AtTKPR2      CAKSKATLKRIVLTSSCSSIRYR--FDATEASP-LNESHWSDPEYCKRFN---LWYGYAK 164
SmTKPR1      CAKS-PSVRRVVLTSSTSAIRFMPEMPSN---SVLDDTSWSSEDFCRKYK---MWYYLAK 172
PpTKPR1      CAKS-TTLKRVVLTSSTAAVRFRDDLEQPGAVTYLDEYSWSSIFFCTKYQ---IWYSLAK 173
Mk657        VKATGGVRRVVL-TSSVAAIWWFA---DRDPEAQLDESYWSDEEYCRENK---LWYQLSK 166
Zc6981       VAATDGIRRVVL-TSSTAAVAWDP---NRDPTAQLDESCWSSEEFCIQNK---MWYHLSK 168


AtANR        VLAEKTAWEFAKENKINLVTVIPALIAGNSLLSDPPSSLSLSMSFITGKEMHVTGLKEMQ 233
SbANR        VLLEKAACRFAEEHGISLVTVCPVLTVGAAPAPKVRTSIVDSLSMLSGDEAGLAVLRGIE 235
PtDFR        TLAEQAAWKFAKENNLDFISIIPTLVVGPFIMQSMPPSLLTALSLITGNEAHYGILKQGH 227
AtDFR        TLAEKAAWDFAEEKGLDFISIIPTLVVGPFITTSMPPSLITALSPITRNEAHYSIIRQGQ 227
PvDFR        TLADKAAIAYAAEHGMDLISVIPPLVIGPFISAGMPPSLLTALALITGNEPHYSILKQVQ 242
AtCCRL14     TLAEDAAWKLAKEKGLDIVTINPAMVIGPLLQPTLNTSAAAILNLING-----AKTFPNL 224
MtCCR        TVAEQSAWDIAKENQVDLVVVNPVVVLGPLLQPTINASTIHILKYLNGA----AKTYVNA 225
AtCCR1       MVAEQAAWETAKEKGVDLVVLNPVLVLGPPLQPTINASLYHVLKYLTGS----AKTYANL 221
PpCCR        LIAERTAWDYADAHGMKLVTICPPVTLGTMLQPRVNQSSKHILKYLDGS----AKTYANR 332
AaTKPR2      TLAEQQAWEFARAHNVDLVVVNPSFVIGPMLPPMPTSTILVVLHLLRG----GAKTFANQ 165
GbTKPR       TIAEKEAWKFAEEKGVNLVVVNPSCVIGPLLAPAPTSTLGIVLSIMRGE---NNGVYPNW 220
Os01g03670   TLAEREAWRLAKERGLDMVAVNPSFVVGPILSQAPTSTALIVLALLRG----ELPRYPNT 232
AtTKPR2      TLGEREAWRIAEEKGLDLVVVNPSFVVGPLLGPKPTSTLLMILAIAKG----LAGEYPNF 220
SmTKPR1      TVAERKAWEFAEKNNLDLVTVLPSFVVGPVLPKNLSSTALDVLGLLKGTVCDVKKFSIYP 232
PpTKPR1      ILSEQEAWKFAFLHSIDLVVVLPSFVIGPCLPYPLSKTAQDICDLLNG----GRNFGIHG 229
Mk657        TQAERAAWAFARESGIELVCINPAMVVGKLLQPSMNTSSEAILKYVTGE----NTVYPNA 222
Zc6981       TQAEKAAWKFAEEKGIDLVTINPAMVIGKMLQPTLNTSSASILKYLTGE----NQTYPNS 224
```

```
AtANR         KLSGSISFVHVDDLARAHLFLAEKETASGRYICCAYNTSVPEIADFLIQRYPKYNVLSEF 293
SbANR         TTSGALQLVHIDDLCRAELFLAEEAAAGGRYICCSLNTTVVELARFLAHKYPQYRVKTNF 295
PtDFR         -------YVHLDDLCMSHIFLYENPKAEGRYICNSDDANIHDLAKLLREKYPEYNVPAKF 280
AtDFR         -------YVHLDDLCNAHIFLYEQAAAKGRYICSSHDATILTISKFLRPKYPEYNVPSTF 280
PvDRF         -------FVHLDDLCDAEIFLFEHPAAAGRYVCSSHATTIHGLAAMLRERYPEYRIPERF 295
AtCCRL14      SFG----WVNVKDVANAHIQAFEVPSANGRYCLVERVVHHSEIVNILRELYPNLPLPERC 280
AtCCR1        TQA----YVDVRDVALAHVLVYEAPSASGRYLLAESARHRGEVVEILAKLFPEYPLPTKC 277
PpCCR         CQA----YVDVKNAAEAHVLAFESPAASGRYLCCKWSLHRGEIVEALARMYPQYAISMRC 388
AaTKPR2       IIG----FVHIDDVVTAHLLVYEEPTASGRYICSERVAHWAEVIKMLKTQTPDYPLPLSP 221
GbTKPR        RLG----FVHIDDAITAHILAMEVPSASGRIICSGDVAHWGEIVEMLRKKYPMYPIADRC 276
Os01g03670    TVG----FVHVDDAVLAHVVAMEDARASGRLICSCHVAHWSEIVGSLRERYPGYPIPAEC 288
AtTKPR2       TVG----FVHIDDVVAAHVLAMEEPKASGRIICSSSVAHWSEIIELMRNKYPNYPFENKC 276
SmTKPR1       RMG----YVHVEDVALAHILVMEAPGARGRYICSSTVMDNDKLGELLAMRYPQFNVP-TK 287
PpTKPR1       RMG----YVHVDDVARAHILVYETPSAQGRYICSAQEATPQELVQYLADRYPHLQIS-TK 284
Mk657         SVG----WVHVEDVAEAHILAYEKPEAEGRYMCVGSWVHWKEITGWLRHLEPSAPITDKC 278
Zc6981        VMG----WVHVEDVADAHILAYENPKAEGRYLCVCTYLHWRELLTALQQICPEAHITQKC 280


AtANR         EEGLS-IPK—LTLSSQKLINEGFRFEYGINEMYDQMIEYFESKGLIKAK          340
SbANR         DDDEHLLERPRVIMSSEKLVREGFEYRHNTLDEIYDNVVEYGKALGILPY         345
PtDFR         KDI-DENLAC-VAFSSKKLTDLGFEFKYSLEDMFAGAVETCREKGLIPLSHRKQVVEE 336
AtDFR         EGV-DENLKS-IEFSSKKLTDMGFNFKYSLEEMFIESIETCRQKGFLPVSLSYQSISE 336
PvDFR         RGI-DDGDLQPVHFSSKKLLDLGFAFKYTVEDMYDAAIRTCREKGLIPLATAGGDGPG 352
AtCCRL14      VD-ENPYVPT-YQVSKDKTRSLGID-YIPLKVSIKETVESLKEKGFAQF          326
AtCCR1        KDEKNPRAKP-YKFTNQKIKDLGLE-FTSTKQSLYDTVKSLQEKGHLAPPP        326
PpCCR         KDDGQPRRVP-LRFCSDKVEQLGLQ-FTSFDETLRNAVSSLQAKGMLHKKTNILKLGS 444
AaTKPR2       NLEEKGNEIP-HRLNTGKIERLGLALFKSLETMFQDCINSFKQN               264
GbTKPR        GVEQ-GNDTP-HTMDTSKIRSLGLSSFKSTEKMFEDCIKSFQEKGLLEGLPLPDHHKH 332
Os01g03670    GSHK-GDDRA-HKMDTAKIRALGFPPFLSVQQMFDDCIKSFQDKGLLPPHA        337
AtTKPR2       SNKE-GDNSP-HSMDTRKIHELGFGSFKSLPEMFDDCIISFQKKGLL            321
SmTKPR1       FPESY-KSKY-YTLDTSKLEKLGLK-FRSVEDMFDDCLENFYHRGLFSLTDTLY     338
PpTKPR1       FNDELPKMPY-YKLNTTKLQRLGLN-CKPLDVMFDDCISFLEEKGLLKRKPEKTPTSS 340
Mk657         TAEDQELVIP-PLMSNEKMQKLGLE-FKSIETMLRDCVDSLKEKNFVKLESSA      329
Zc6981        AD-DNPDALP-PLLNTDKLKGLGLH-FRTMQTMLIDTAASLRERGFYKP          326
```


[1] Petit P, Granier T, d'Estaintot BL, Manigand C, Bathany K, Schmitter J-M, Lauvergeat V, Hamdi S, Gallois B. 2007. Crystal structure of grape dihydroflavonol 4-reductase, a key enzyme in flavonoid biosynthesis. *J Mol Biol* **368**: 1345–1357.

[2] Lewis JA, Zhang B, Harza R, Palmer N, Sarath G, Sattler SE, Twigg P, Vermerris W, Kang C. 2023. Structural similarities and overlapping activities among dihydroflavonol 4-reductase, flavanone 4-reductase, and anthocyanidin reductase offer metabolic flexibility in the flavonoid pathway. *Int J Mol Sci* **24**: 13901.

[3] Chao N, Li S, Li N, Qi Q, Jiang W-T, Jiang X-N, Gai Y. 2017. Two distinct cinnamoyl-CoA reductases in *Selaginella moellendorffii* offer insight into the divergence of CCRs in plants. *Planta* **246**: 33–43.

[4] Sattler SA, Walker AM, Vermerris W, Sattler SE, Kang C. 2017. Structural and biochemical characterization of cinnamoyl-CoA reductases. *Plant Physiol* **173**: 1031–1044.
